# New GFAP splice isoform (GFAPμ) differentially expressed in glioma translates into 21 kDa N-terminal GFAP protein

**DOI:** 10.1101/2020.07.04.187443

**Authors:** Emma J. van Bodegraven, Jacqueline A. Sluijs, A. Katherine Tan, Pierre A.J.T. Robe, Elly M. Hol

## Abstract

The glial fibrillary acidic protein (GFAP) is a type III intermediate filament (IF) protein that is highly expressed in astrocytes, neural stem cells, and in gliomas. Gliomas are a heterogeneous group of primary brain tumors that arise from glia cells or neural stem cells and rely on accurate diagnosis for prognosis and treatment strategies. GFAP is differentially expressed between glioma subtypes and therefore often used as a diagnostic marker. However, GFAP is highly regulated by the process of alternative splicing; many different isoforms have been identified. Differential expression of GFAP isoforms between glioma subtypes suggests that GFAP isoform-specific analyses could benefit diagnostics. In this study we report on the differential expression of a new GFAP isoform between glioma subtypes, GFAPμ. A short GFAP transcript resulting from GFAP exon 2 skipping was detected by RNA sequencing of human glioma. We show that GFAPμ mRNA is expressed in healthy brain tissue, glioma cell lines, and primary glioma cells and that it translates into a ^~^21 kDa GFAP protein. 21 kDa GFAP protein was detected in the IF protein fraction isolated from human spinal cord as well. We further show that induced GFAPμ expression disrupts the GFAP IF network. The characterization of this new GFAP isoform adds on to the numerous previously identified GFAP splice isoforms. It emphasizes the importance of studying the contribution of IF splice variants to specialized functions of the IF network and to glioma diagnostics.

**Summary statement:** RNA sequencing data of glioma patient material reveals the differential expression of a short GFAP splice isoform, *GFAPμ*, between glioma subtypes. *GFAPμ* translates into a 21 kDa N-terminal GFAP protein which can disrupt the GFAP intermediate filament network.

## Introduction

Glial fibrillary acidic protein (GFAP) is a type III intermediate filament (IF) protein, which forms together with vimentin, synemin, and nestin the cytoskeletal IF-network in astrocytes. GFAP is often used as a marker of astrocytes, and increased expression of GFAP as an indicator of reactive gliosis in the injured and diseased brain (Middeldorp and Hol, 2011; Hol and Pekny, 2015). Moreover, GFAP is a classical diagnostic marker for the most malignant tumors of the central nervous system (CNS); gliomas (Velasco et al., 1980; Dunbar and Yachnis, 2010; Perry and Wesseling, 2016). Gliomas are a large and diverse group of primary brain tumors that arise from glia cells or their precursors, and GFAP is differentially expressed between glioma subtypes (van Bodegraven et al., 2019a).

However, the GFAP gene is highly regulated by the process of alternative splicing (Blechingberg et al., 2007b; Kanski et al., 2014). Up to now, the expression of 8 different human GFAP transcripts has been confirmed; the canonical variant GFAPα (Reeves et al., 1989), GFAPδ (Nielsen et al., 2002; Roelofs et al., 2005), GFAPγ (Zelenika et al., 1995), GFAPκ (Blechingberg et al., 2007a), GFAPΔ135 (Hol et al. 2003), GFAPΔ164 (Hol et al. 2003), GFAPΔexon6 (Hol et al. 2003), and recently GFAPλ (Helman et al., 2020). These GFAP isoforms are expressed in specific cell types of the CNS, such as radial glia and adult neural stem cells, in reactive gliosis in Alzheimer’s disease and epilepsy, in Alexanders disease, and in glioma (Middeldorp and Hol, 2011; Hol and Capetanaki, 2017; Helman et al., 2020). The isoforms are generated by the in- or exclusion of an intronic or exonic region that changes the coding sequence (CDS) of the transcript, the 5’ untranslated region (UTR), and/or the 3’ UTR.

In recent years, higher resolution methods such as RNA sequencing have shown that the level of RNA processing varies between tissue types, developmental stages, and between healthy and diseased tissues (Ameur et al., 2011; Elkon et al., 2013; Bentley, 2014; Herzel et al., 2017). Altered RNA splicing (Dvinge and Bradley, 2015; Lee and Abdel-Wahab, 2016) and increased 3’UTR shortening (Morris et al., 2012) is observed in various cancer types. For example, alternative splicing events that result in premature termination codons (PTC) are more often observed in cancer compared to healthy tissue (Chen, Tovar-Corona, and Urrutia 2011). Expression of transcripts that contain a PTC, but that are not degraded by the mechanism of nonsense-mediated RNA decay, results in the translation of shorter proteins (Hug et al., 2016). As changes in RNA processing can function as a pro-oncogenic mechanism (Fu et al., 2011), targeting RNA processing to treat cancer is currently under investigation (Lee and Abdel-Wahab, 2016).

In our previous studies, we have reported on alterations in GFAP alternative splicing in glioma. We have observed that in high malignant glioma the relative level of the alternative splice variant GFAPδ to the canonical variant GFAPα is increased compared to lower malignant glioma (Stassen et al., 2017) and *in vitro* studies suggest that changes in the relative level of GFAPδ to GFAPα have functional implications for glioma (Moeton et al., 2014; Stassen et al., 2017; van Bodegraven et al., 2019b). These studies emphasize the importance of discriminating between GFAP splice variants when studying the role of GFAP in glioma malignancy and using GFAP as a diagnostic marker (van Bodegraven et al., 2019a).

We here report on the differential expression of a new GFAP isoform in glioma of different grades of malignancy, GFAPμ. Based on the annotated sequence in the RNA sequencing data of human glioma (The Cancer Genome Atlas) we identified GFAPμ as a product of skipping of GFAP exon 2. This leads to the generation of a transcript with a PTC in exon 3 and consequently an extremely short CDS. We confirm the endogenous expression of GFAPμ mRNA in different types of tissue and cells and show translation of GFAPμ cDNA into a ^~^21 kDa sized GFAP protein that is detected in the human spinal cord as well. We further characterize GFAPμ upon its induced expression in different cell lines and describe how its expression influences the IF network.

## Material and Methods

### RNA sequencing TCGA data analysis

RNA sequencing data of 165 grade IV glioma and 306 low grade glioma (including glioma of the astrocytoma histological subtype) was obtained from the cancer genome atlas (TCGA). Normalized RNA isoform expression data (Level 3 released data downloaded June 2015) was extracted as upper quantile normalized RSEM (RNA-Seq by Expectation Maximization) count estimates (normalized expression). Recurrent tumors were removed, and normalized counts of duplicate tumor samples were averaged. Isocitrate dehydrogenase 1 (IDH1) and short arm of chromosome 1/long arm of chromosome 19 (1p/19q) status information was available for 144 grade IV glioma and 282 low grade glioma. For grade IV glioma, processed broad mutation data was downloaded from the UCSC Cancer Browser in June 2015. For low grade glioma data was extracted from the TCGA network publication of 2015 (The Cancer Genome Atlas Research Network, 2015). Glioma were categorized according to the World Health Organization classification system of 2007 (histological subtypes) or 2016 (molecular subtypes). Supplementary Table 1 contains information on the included patient cohort.

### Sample collection

Fresh frozen healthy temporal cortex tissue from three different donors was provided by the Netherlands Brain Bank (NBB; hersenbank.nl). Tissue was lysed using TRIzol reagent (Ambion by Thermo Scientific, Waltham, MA, USA). Primary adult neural stem cells were isolated from post-mortem brain tissue and lysed according to a previously described protocol (van Strien et al., 2014).

Primary glioma cells were obtained from tumor tissue of patients undergoing resection surgery for glioma grade IV at the University Medical Center Utrecht. Tumor tissue samples were placed directly into tissue culture flasks and maintained in DMEM/F-12 (Gibco, Thermo Scientific, Waltham, MA, USA) supplemented with 100 U/ml penicillin, 100 μg/ml streptomycin (1% p/s) and 10% (v/v) Fetal Bovine Serum (FBS) (all Invitrogen, Bleiswijk, The Netherlands). Cells were passaged at confluence for maintenance and collected and lysed using TRIzol reagent (Ambion by Thermo Scientific, Waltham, MA, USA). IF fractions from human spinal cord were obtained as described previously (Perng et al., 2008).

### Cell lines and culture

All cell lines were maintained at 37°C in a humidified incubator with 5% CO2. U251-MG glioma cells (obtained from Muenster, Germany) were maintained in DMEM high glucose: Ham’s F10 nutrient mix, supplemented with 1% p/s and 10% FBS (all Invitrogen, Bleiswijk, The Netherlands). The identity of U251-MG cells was confirmed by short terminal repeat analysis (Eurofin, Luxembourg). Human embryonic kidney 293 cells (293T) were maintained in DMEM high glucose supplemented with 1% p/s and 10 % (v/v) FBS. The adrenal carcinoma cell lines SW13.Vim- and SW13.Vim+ were maintained in DMEM:Ham’s F12 nutrient mix and glutaMAX supplemented with 1% p/s and 5% (v/v) FBS. Human immortalized foetal neural stem cells (ihNSCs) (De Filippis et al., 2007) were maintained in Euromed-N medium (EuroClone, Amsterdam, The Netherlands), 1% N2 (5.375 mL DMEM-F12 (Gibco, Thermo Scientific, Waltham, MA, USA), 0.75% BSA, 62.5 mg insulin (Sigma, Saint-Louis, MO, USA), 100 mg apo-transferrin (Sigma), 10 μL 3 mM Na-Selenite (Sigma), 16 mg Putrescine (Sigma), 20 μg Progesterone (Sigma)), 1% glutaMAX (Gibco), 1% L-glutamin (Gibco), 1% p/s, 20 ng/ml EGF and 10 ng/ml FGF (both Tebu-Bio, Heerhugowaard, The Netherlands).

### RNA isolation

For RNA isolation of cell lines, cells were plated at a density of 4 x 10^4^ cells per well in a 24 well plate. After 3 days cells were lysed using TRIzol (Ambion by Thermo Scientific, Waltham, MA, USA). To extract RNA from lysed tissue and cell line samples in TRIzol, chloroform (EMD Millipore Inc., Darmstadt, Germany) was added and by centrifugation at 12,000 g at 7°C for 15 min., RNA was separated from proteins and lipids. RNA was precipitated in 2-propanol (EMD Millipore Inc., Darmstadt, Germany) at −20°C overnight and centrifugation at 16,000 g at 4°C for 45 min. resulted in a RNA containing pellet. Pellets were washed twice with 75% cold ethanol and dissolved in MilliQ. RNA concentrations and purity were measured using the Varioskan Flash (Thermo Scientific, Waltham, MA, USA).

### cDNA synthesis and real-time quantitative PCR analysis

To generate cDNA, the Quantitect Reverse Transcription kit (Qiagen, Hilden, Germany) was used according to the manufacturer’s protocol. In short, DNAse (gDNA wipe out buffer, Qiagen) was added to ^~^500 ng RNA and activated at 42°C for 2 min. The RNA was converted to cDNA in a 10 μl reaction mix that contained reverse transcriptase enzyme, reverse transcriptase buffer, random primers (hexanucleotides), and oligo-dTs (all Qiagen) at 42°C for 30 min. followed by 3 min. at 95°C. cDNA was diluted 10x in MilliQ and 1 μl was used for real time qPCR analysis in a mix containing 1 μl of primer mix (final concentration of 0.1 μM for forward and reverse primer), 5 μl FastStart Universal SYBR Green Master mix (ROX) (Roche, Basel, Switzerland) and 3 μl MilliQ. The reaction mix was added to a 96 or 384 plate and amplification of the product was measured after incubation steps at 50°C for 2 min. and 95°C for 10 min., during 40 PCR cycles (95°C for 15 sec. and 60°C for 1 min.) using a QuantStudio 6 Flex Real-Time PCR System (Applied Biosystems, Foster City, CA, USA). A dissociation curve was generated afterwards by ramping the temperature from 60°C to 95°C to determine the specificity of the PCR product. For detection of the GFAP isoforms the following primers were used: pan GFAP forward primer 5’-GACCTGGCCACTGTGAGG-3’, reverse primer 5’- GGCTTCATCTGCTTCCTGTC-3’, GFAPα forward primer 5’-TAGGCTCTCTCTGCTCGGTT-3’, reverse primer 5’-GAGGGCGATGTAGTAGGTGC-3’, and GFAPμ forward primer 5’-TGGCCACTGTGAGGCAGAAGAAG-3’, reverse primer 5’- TCATGCATGTTGCTGGACGC-3’. To determine specific amplification of the GFAPμ products of the qPCR, products were separated by electrophoresis in an 8% acrylamide gel made in TBE Buffer (18 mM tris (hydroxymethyl)-aminomethane, 17.8 mM boric acid, 0.4 mM EDTA pH 8.0 in MilliQ)), stained using SYBR Safe DNA Gel Stain (Invitrogen, Carlsblad, CA, USA) and imaged using an E-Gel Imager System with Blue Light Base (Life Technologies, Carlsblad, CA, USA).

### Plasmid construction

Using two different primer sets, GFAP exon 1 and GFAP exon 3 CDS sequences were generated by PCR from pcDNA3.1 containing full length GFAP cDNA as a template (1 μg plasmid DNA, 10 pmol/μl forward and reverse primer, PFU buffer, PFU enzyme, 10 mM dNTPs; annealing at 62°C, elongation for 5 min.). Products were separated using gel electrophoreses and correct size products were isolated using a gel extraction kit according to the manufacturer’s protocol. Exon 1 was digested at its 5’ using *Hind*III and exon 3 was digested at its 3’ using *Bam*H I (3 hours at 37°C). Exon 1 and exon 3 digestion products were purified using the High pure PCR product purification kit (Roche, Basel, Switzerland) according to the manufacturer’s protocol. An overnight blunt ligation was induced for the 5’ of exon 1 and the 3’ of exon 3 (5 U T4 DNA ligase (Roche, Basel, Switzerland)) at room temperature. *Hin*dIII and *Bam*HI were again used to digest pcDNA3.1 (3 hours at 37°C) and the digestion product was separated using gel electrophoreses, isolated, and purified as described above. Exon 1 and exon 3 were ligated in pcDNA3.1 overnight at room temperature. The correct sequence was verified by sequencing (Macrogen, Amsterdam). To generate plasmid DNA containing GFAPμ cDNA and an upstream sequence encoding 20 amino acids that is recognized by the biotinylation enzyme BirA, pcDNA3.1-GFAPμ was digested using HindIII. The double stranded biotin-tag oligo was digested with HindIII as well and ligated into the pcDNA3.1-GFAPμ plasmid. The orientation of the biotin-tag was verified by sequencing (Macrogen, Amsterdam).

### Transfection

For immunostaining, cells were plated on laminin coated coverslips in a 24 well plate at a density of 2 x 10^4^ cells per well (SW13.Vim-, and SW13.Vim+ cells) or 5 x 10^3^ cells per well (U251-MG cells). For western blot analysis, cells were plated in a 6 well plate at a density of 5 x 10^5^ cells per well (293T cells). Cells were transfected with empty pcDNA3.1 (mock), pcDNA3.1-GFP (GFP), pcDNA3.1-GFAPμ (GFAPμ), pcDNA3.1-bio-GFAPμ (bio-GFAPμ), pcDNA3.1-bio-GFAPα (bio-GFAPα), and/or pCI-neo-BirA (BirA) using polyethyleneimine (166 ng/ml final concentration). Expression of plasmid DNA was allowed for 3 to 5 days depending on the experiment before cells were fixed in 4% paraformaldehyde (PFA) in phosphate buffer saline (PBS, pH 7.2) for immunostaining or cell pellets were harvested for western blot analysis.

### Immunocytochemistry

Coverslips with cells were incubated in blocking buffer (50 mM Tris pH 7.4, 150 mM NaCl, 0.25% (w/v) gelatin, and 0.5% triton X-100) at room temperature for 15 min., followed by an overnight incubation with the primary antibody in blocking buffer at 4°C. Coverslips were washed in PBS, incubated in secondary antibody in blocking buffer at room temperature for 1 hour, washed again in PBS, dipped in MilliQ, and mounted in Mowiol (0.1 M tris-HCl pH 8.5, 25% glycerol, 10% Mowiol (Calbiochem, Merck Millipore, Darmstadt, Germany)). To counterstain the nuclei of the cells, Hoechst (1:1000, 33528, Thermo Fisher Scientific, Waltham, MA, USA) was used and co-incubated with secondary antibodies. To label biotinylated proteins, fluorescently labelled streptavidin (1:1,400 Alexa Fluor 488, Jackson Immuno Research, West Grove, PA, USA) was co-incubated. Immunofluorescent images were taken using a Zeiss Axioscope.A1 microscope. Table 1 provides a list of antibodies used in this study.

**Table 1.** List of antibodies.

| **Antibody** | **Product Code, Company** | **Dilution Western Blot** | **Dilution Immunocytochemistry** |
| --- | --- | --- | --- |
| Rabbit anti-GFAP (panGFAP) | #Z0334, Dako (Agilent), Santa Clara, CA, USA | 1:50,000 | 1:4,000 |
| Goat anti-GFAP N-18 (N-term) | sc-6171, Santa Cruz, Dallas, Texas, USA | 1:1,000 | 1:1,000 |
| Chicken anti-Vimentin | AB5733, Chemicon, Temecula, California | - | 1:4,000 |
| Donkey anti-rabbit Cy3 | Jackson Immuno Research, West Grove, PA, USA | - | 1:1,000 |
| Donkey anti-goat 488 | Jackson Immuno Research, West Grove, PA, USA | - | 1:1,000 |
| Donkey anti-rabbit IRdye800 | Jackson Immuno Research, West Grove, PA, USA | 1:5,000 | - |
| Donkey anti-goat AF647 | Jackson Immuno Research, West Grove, PA, USA | 1:2,000 | - |
| Donkey anti-chicken Dylight488 | Jackson Immuno Research, West Grove, PA, USA | 1:2,000 | - |

### Western Blot analysis

Cell pellets were resuspended in suspension buffer (0.1 M NaCl, 0.01 M Tris-HCl pH 7.6, 0.001 M EDTA, cOmplete EDTA-free protease inhibitor cocktail (Roche, Basel, Switzerland)) and lysed in 2x SDS loading buffer (100 mM Tris pH 6.8, 4% SDS, 20% glycerol, 5% 2-mercaptoethanol, bromophenol blue).

Samples were heated at 95 °C for 5 min. after which the DNA was sheared using a 25 gauge needle. Samples were loaded on a 15% SDS-PAGE gel and proteins were separated by electrophoresis. Proteins were blotted on a 0.45 μM pore size nitrocellulose membrane (GE Healthcare, Chicago, IL, USA) using a Transblot SD semi-dry transfer system (Bio-rad, Hercules, CA, USA) system for 1 hour. Blots were incubated in blocking buffer (50 mM Tris pH 7.4, 150 mM NaCl, 0.25% (w/v) gelatine, and 0.5% triton X-100) and incubated in primary antibody in blocking buffer at 4°C O/N. Blots were washed 3 times in TBS-T (100 mM Tris-HCl pH 7.4, 150 mM NaCl and 0.2% Tween-20) and incubated in secondary antibody in blocking buffer at room temperature for 1 hour. To label biotinylated proteins, blots were incubated with fluorescently labelled streptavidin (1:2,000 Alexa Fluor Cy5, Jackson Immuno Research, West Grove, PA, USA). Blots were washed 3 times in TBS-T and ones in milliQ before blots were scanned using the Odyssey CLx Western Blot Detection System (LI-COR Biosciences, Lincoln, NE, USA). Table 1 provides a list of used antibodies.

### Statistics

To determine differential expression of GFAPμ the data was tested for a normal distribution and normality of variances using the Shapiro Wilk test and Levene’s test. A one-way ANOVA or Kruskal-Wallis test was performed dependent on the distribution of the data followed by a post-hoc Tuckey’s honestly significant difference (HSD) tests or Nemenyi tests, respectively. All analyses were performed using R software (version 3.4.3) and the PMCMR package (version 4.3). Survival analysis (including progression free survival) were performed using the Survival package (version 2.41-3). Kaplan Meier survival curves were compared using a log-rank regression analysis.

## Results

### Differential expression of a short GFAP transcript in glioma subtypes

The analysis of RNA sequencing data of glioma patients from The Cancer Genome Atlas (TCGA)(Supplementary Material 1) revealed the expression of a third GFAP isoform in addition to the well-known GFAPα and GFAPδ isoforms (Stassen et al., 2017). The CDS of this third GFAP isoform, GFAPμ, consisted of only exon 1 and exon 3 of the GFAP gene according to the RNA sequencing transcript annotation file (Supplementary Material 2) and the Ensembl database (GFAP-202, www.ensembl.org). GFAPμ was expressed in astrocytoma grade II, III, and IV (Figure 1A) and the expression was significantly decreased in grade IV astrocytoma patients. This expression pattern follows the decrease of the canonical GFAPα isoform in high grade astrocytoma as we previously reported (Stassen et al., 2017). However, the overall expression level of GFAPμ was about 5,000 times lower. The GFAPμ expression levels did not correlate with patient prognosis within grade II, III or IV astrocytoma (data not shown).

**Figure 1.**
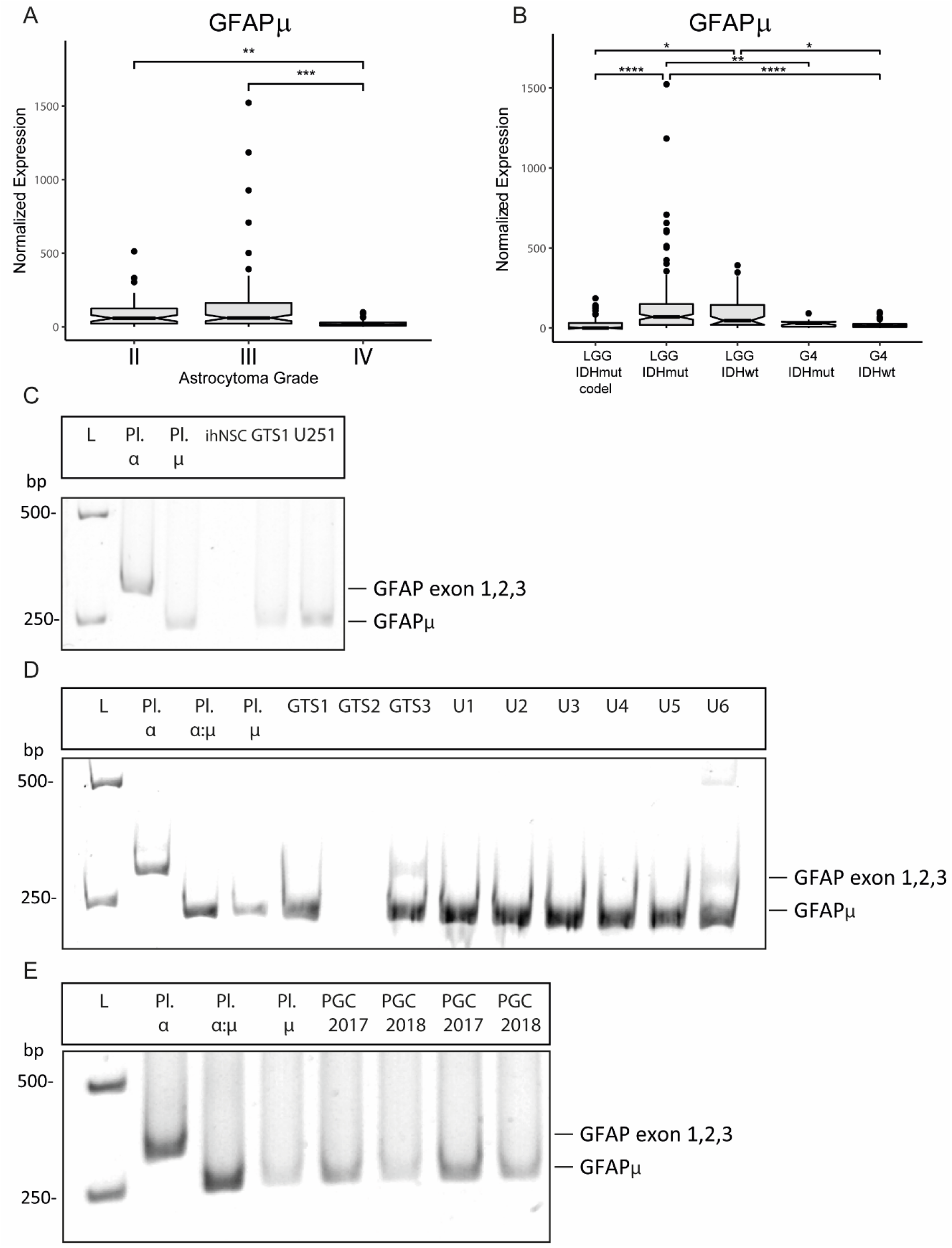
GFAPμ expression in glioma subtypes, glioma primary cells and cell lines, and in healthy brain tissue. **A, B** Normalized expression of a new GFAP isoform (GFAPμ) in astrocytoma of grade II, III and IV (WHO 2007) and in different glioma subtypes (WHO2016). Expression levels were obtained from RNAseq level 3 released normalized isoform expression data of the TCGA database (*LGG* = grade II and III glioma, *G4* = grade IV glioma, *IDHmut* = IDH1 mutation, *codel* = 1q19p co-deletion, *whiskers*: ±1.5 x IQR; *notch:* 95% CI). **C, D, E** Images of qPCR products separated on an 8% acrylamide gel by electrophoreses. 252 bp sized products are generated from GFAPμ transcripts. 313 bp sized products are generated from GFAP transcripts that contain exon 1, 2 and 3 (*L* = ladder, *Pl. α* = GFAPα plasmid cDNA, *Pl. μ* = GFAPμ plasmid cDNA, *ihNSC* = immortalized adult human neural stem cells, *GTS1-3* = 3 different human temporal cortex samples, *U251* = U251-MG glioma cells, *U1-U6* = different U251-MG clonal cell lines, PGC = primary glioma cells isolated in 2017 and 2018). *\* p < 0.05, ** p < 0.01, *** p < 0.001, **** p < 0.0001*

Recently, the World Health Organization has changed the glioma classification system (WHO 2016) based on additional molecular tumor characteristics. Expression of GFAPμ in these glioma subtypes is shown in Figure 1B. GFAPμ was higher expressed in low-grade IDH1 mutated and IDH1 wild type tumors without a 1p/19q deletion compared to grade IV IDH1 mutated and wild type tumors. GFAPμ expression in IDH1 mutated tumors with a 1p/19q deletion was lower compared to low grade IDH1 mutated and wild type tumors as well. Within these glioma subtypes, GFAPμ did not have a significant prognostic value for patients either (data not shown).

### GFAPμ mRNA expression in human brain tissue and glioma cells

Generation of the GFAPμ transcript must be proceeded by a new alternative splicing event of the GFAP gene that leads to skipping of exon 2. Skipping of exon 2 induces a frame-shift and PTC in exon 3 of the resulting transcript (Supplementary Material 2 and 3). The CDS of GFAPμ therefore only consists of 540 base pairs (bp), which is less than half of the 1299 bp CDS of GFAPα (Figure 2B, Supplementary Material 2). To determine if the skipping of GFAP exon 2 is an alternative splicing event that is observed more often, we performed qPCR analysis on different samples using GFAPμ specific primers followed by gel electrophoresis (Figure 1C-E). The forward primer was designed to bind to the 3’ end of exon 1 and two nucleotides at the 5’ end of exon 3 (thus spanning exon 1 and 3). The reverse primer was designed to bind exon 3 downstream of the forward primer to generate a 252 bp product. Figure 1C lane 2 shows that this primer pair could also amplify GFAPα cDNA that contains exon 1, 2 and 3. However, the size of the product is longer (331 bp) as it includes exon 2. Amplification of GFAPμ generated a 252 bp product (Figure 1C lane 3). Figure 1C, lane 5 and 6 show that a 252 bp GFAPμ product was detected in RNA isolated from both healthy human brain tissue and from the human U251-MG glioma cell line. Figure 1D lane 3 and Figure 1E lane 3 show that when combining GFAPα cDNA and GFAPμ cDNA, the amplification of the 252 bp GFAPμ product was preferred. GFAPμ was detected in 2 out of 3 human brain samples (Figure 1D, lane 5-7), and in different U251-MG glioma clonal cell lines (Figure 1D, lane 8-13). In addition, GFAPμ was detected in two primary glioma cell cultures (Figure 1E, lane 5-8). GFAPμ was not detected in ihNSCs (Figure 1C) or adult neural stem cells isolated from human post-mortem tissue that did express pan GFAP (data not shown). These results suggest that the skipping of exon 2 is a common GFAP alternative splicing event that generates stable GFAPμ mRNA molecules in glioma and in the healthy brain.

**Figure 2.**
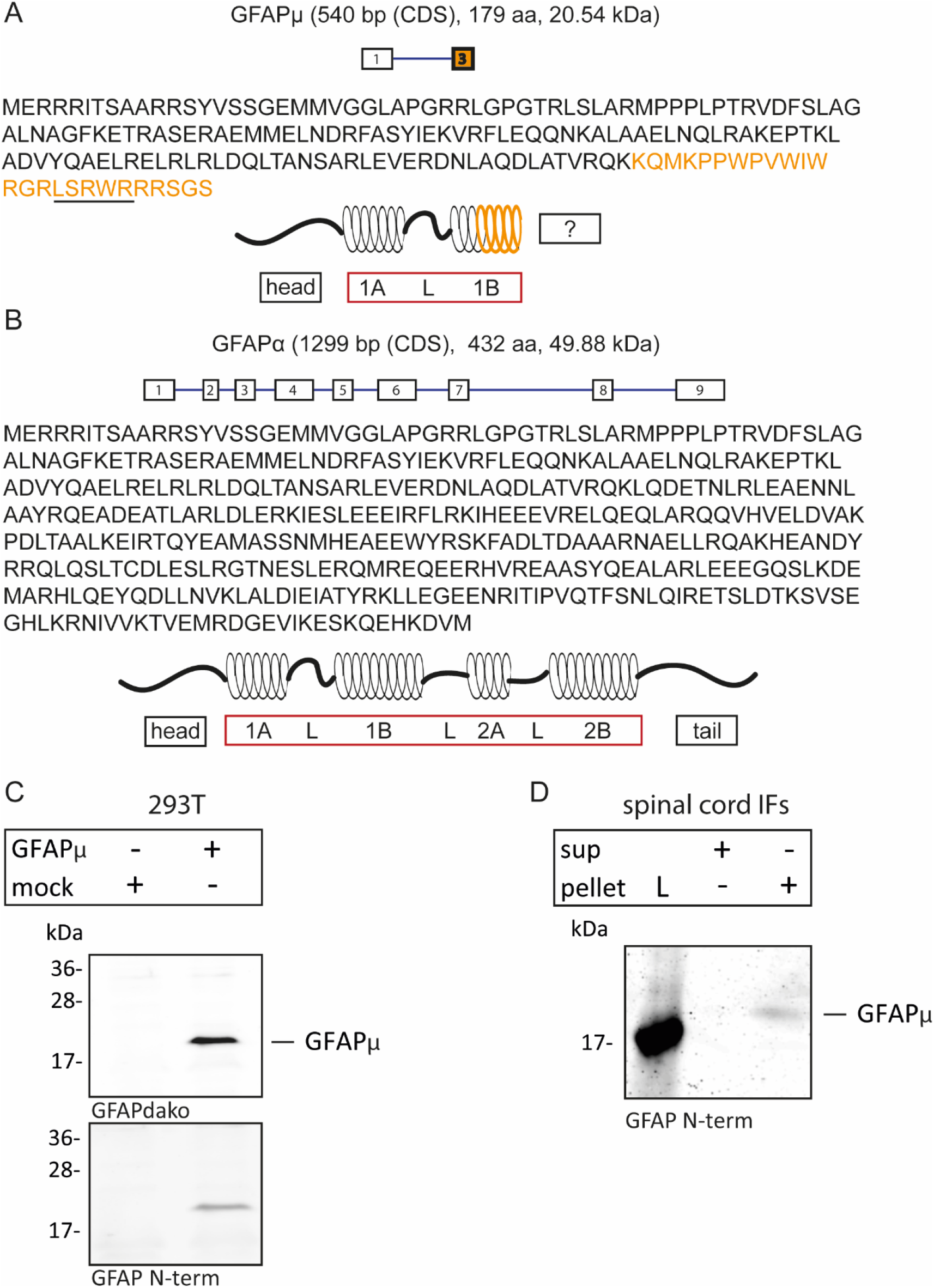
GFAPμ characteristics and western blot analysis. **A, B** Characteristics of GFAPμ and the canonical GFAPα isoform transcripts and proteins and a hypothesized protein structure of GFAPμ (*1A = helical coiled-coiled segment 1A, L = Linker segments, 1B = helical coiled-coiled segment 1, 2A = helical coiled-coiled segment 2A, 2B = helical coiled-coiled segment 2B*). The underlined sequence indicates the peptide that was found in a human proteomics study (Kim et al., 2014). **C** Western blot analysis of whole lysates of GFAPμ and mock transfected 293T cells. Western blots were stained with two different GFAP antibodies that recognize a protein sequence encoded by exon 1 (top) and the N-terminal head of GFAP (bottom). **D** Western blot of intermediate filament protein isolated from the human spinal cord. The blot was stained with a GFAP antibody that recognizes the N-terminal head of the GFAP protein (*L = protein ladder*).

### GFAPμ is translated into a 21 kDa sized protein

The annotation of a unique peptide (Figure 2A, underlined sequence) that specifically aligns to the GFAPμ protein sequence was found in a human proteomics study (Kim et al., 2014) and suggested that GFAPμ is a protein coding alternative splice variant. The data of this study is deposited in a protein database (proteomicsdb.org) where GFAPμ is annotated as GFAP-B1DIR4. Comparison of the canonical isoform GFAPα (Figure 2B) and GFAPμ (Figure 2A) protein sequences showed that the predicted GFAPμ protein lacks a large part of the IF rod domain and its entire tail. To confirm translation of the GFAPμ CDS and stable expression of the protein, we expressed GFAPμ cDNA (Supplementary Material 2) in 293T cells. Western blotting of proteins isolated from GFAPμ and mock transfected cells (Figure 2C) confirmed the expression of a ^~^21 kDa protein recognized by two different GFAP antibodies. Both antibodies recognize the GFAP N-terminal region that is present in GFAPμ. In 293T cells, which have undetectable endogenous GFAP expression, a clear and strong GFAPμ band is visible in the transfected cells (Figure 2C). This confirms that GFAPμ cDNA can translate into a stable protein. Interestingly, on a western blot of IF proteins isolated from the human spinal cord, a ^~^21 kDa size protein was detected by the GFAP N-terminal antibody (Figure 2D), suggesting translation and stable protein expression of the endogenous GFAPμ transcript.

### Limited GFAPμ self-assembly and de novo filament formation

We continued to investigate the behavior of the GFAPμ protein in different cellular environments. GFAP variants lacking essential domains of the N-terminal head, rod or C-terminal tail are assembly compromised and form aggregates in the absence of other IF proteins with which they can co-assemble (Roelofs et al. 2005; Moeton et al. 2016; Perng et al. 2008; Chen and Liem 1994; Nielsen and Jorgensen 2004). To determine whether GFAPμ is capable of *de novo* filament formation in the absence of other IF proteins, we used the human adrenal carcinoma-derived cell line SW13 that is devoid of vimentin (SW13.Vim-) and other cytoplasmic IFs. As described in previous studies (Jing et al. 2007; Chen and Liem 1994), the canonical splice variant GFAPα forms a filamentous network in these cells (Figure 3A). Immunostainings of mock and GFAPμ transfected SW13.Vim-cells for two different GFAP antibodies that recognize GFAPμ are shown in Figure 3B-F. In most cells, GFAPμ expression led to diffuse non-filamentous GFAP immunostaining with small and bright GFAP aggregates throughout the cell’s cytoplasm (Figure 3C, D) and/or larger (peri-)nuclear aggregations (Figure 3E). A rare observation of filament formation is shown in Figure 3F where GFAP positive filament-like bundles were seen around the nuclei of these cells. However, as these cells seem to be dividing this might be a cellular state specific staining pattern and no direct evidence of GFAPμ self-assembly. Confocal imaging showed that in some cells the aggregated structures in the cell periphery were short and thick filament-like structures (Figure 3G and H). Furthermore, Figure 3H shows that aggregates did not overlap with Hoechst staining but were present in the peri-nuclear regions and most likely within the nuclear invaginations (Jorgens et al., 2017). These results do not provide evidence for the ability of GFAPμ to self-assemble into filaments.

**Figure 3.**
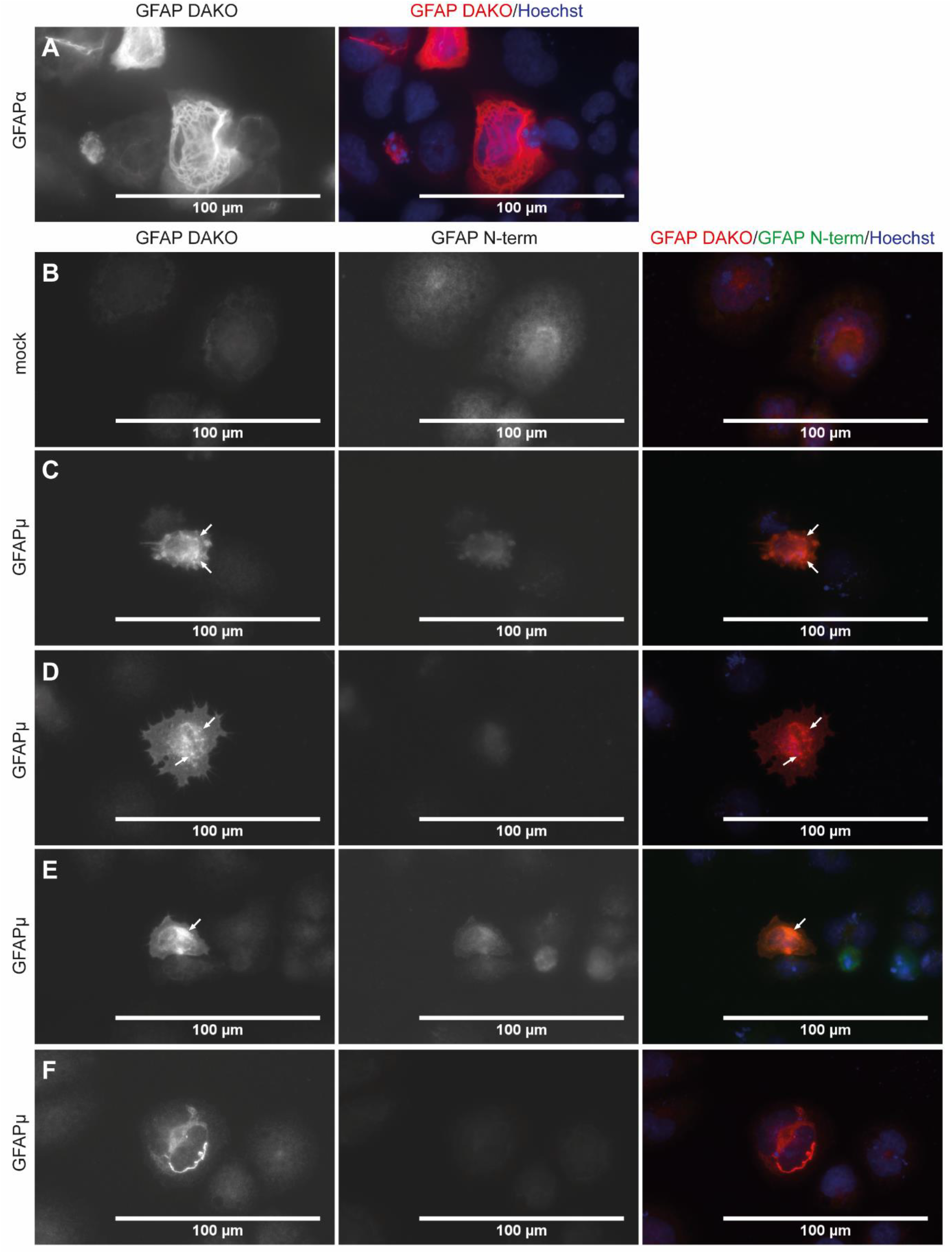
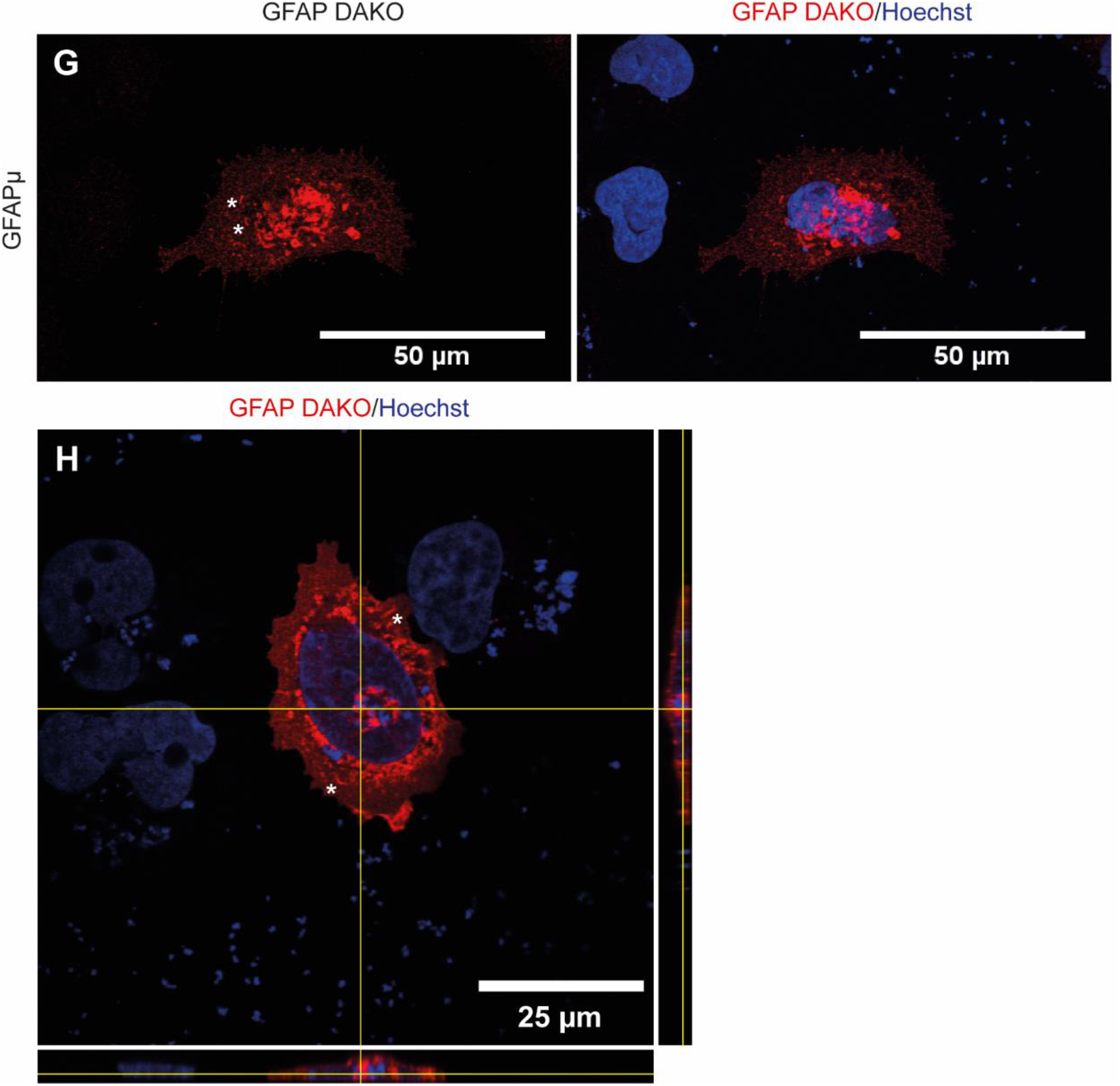
GFAPμ staining patterns in SW13.Vim-cells. Immunostaining images of GFAPα (A), mock (B) and GFAPμ (C-H) transfected SW13.Vim-cells. **A** GFAPα transfected cells stained using a GFAP (Dako) antibody. **B-H** Mock and GFAPμ transfected cells stained using two different GFAP antibodies (GFAP Dako (left), GFAP N-term (middle)) and Hoechst (right). Diffuse GFAPμ expression in small bright and larger peri-nuclear aggregates (arrows indicate aggregates) (C, D). A pattern of diffuse GFAPμ expression and large peri-nuclear aggregation (arrows indicate aggregates) (E). Filamentous expression patterns of GFAPμ around the nuclei (F). Confocal images of thick filamentous structures (G) and peri-nuclear GFAP aggregates (H). Asterisks (*) indicate short and thick filament-like structures.

### GFAPμ expression patterns in the presence of the GFAP assembly partner vimentin

GFAP mutants or isoforms that lack essential domains to self-assemble into filaments can disrupt the existing IF network or make use of IF assembly partners that are present to co-assemble (Roelofs et al. 2005; Moeton et al. 2016; Perng et al. 2008; Chen and Liem 1994; Nielsen and Jorgensen 2004). To determine the behavior of GFAPμ in the presence of the known GFAP assembly partner vimentin, GFAPμ was expressed in SW13.Vim+ cells (Figure 4). Co-immunostaining for vimentin and GFAP showed different patterns of IF expression. In some un-transfected control cells, cross-reactivity of the GFAP Dako antibody with vimentin was observed (Figure 4A), but immunostaining intensity was much stronger and did not co-localize with vimentin in GFAPμ transfected cells (Figure 4B-I). In Figure 4B and C, cells with diffuse GFAP expression are shown that contained brighter aggregated GFAP structures near the nucleus. In these cells, vimentin was mainly present near the nucleus as well and the brighter filamentous structures as seen in most control cells were absent (Figure 4B). However, this vimentin staining pattern was not unique for GFAPμ positive cells and was seen in some un-transfected controls (Figure 4A, B, D) as well. In addition, Figure 4D shows similar diffuse GFAP staining and peri-nuclear aggregates whereas vimentin positive structures were brighter and filamentous. Confocal imaging shows that the GFAP aggregates localize to the intact vimentin network (Figure 4E). The GFAP aggregation observed was similar to the in Figure 3G and H described small thick filamentous structures. Furthermore, some cells contained larger peri-nuclear aggregates of both GFAP and vimentin (Figure 4F). A fourth common pattern is shown in figure 4G and H in which a low cytoplasmic GFAP staining was seen together with bright small GFAP positive aggregates and an intact filamentous vimentin IF network. In some cases, the aggregates lined up along the vimentin network. These results do not provide clear evidence for GFAPμ-induced disruption of the vimentin IF network or vimentin-GFAPμ co-assembly.

**Figure 4.**
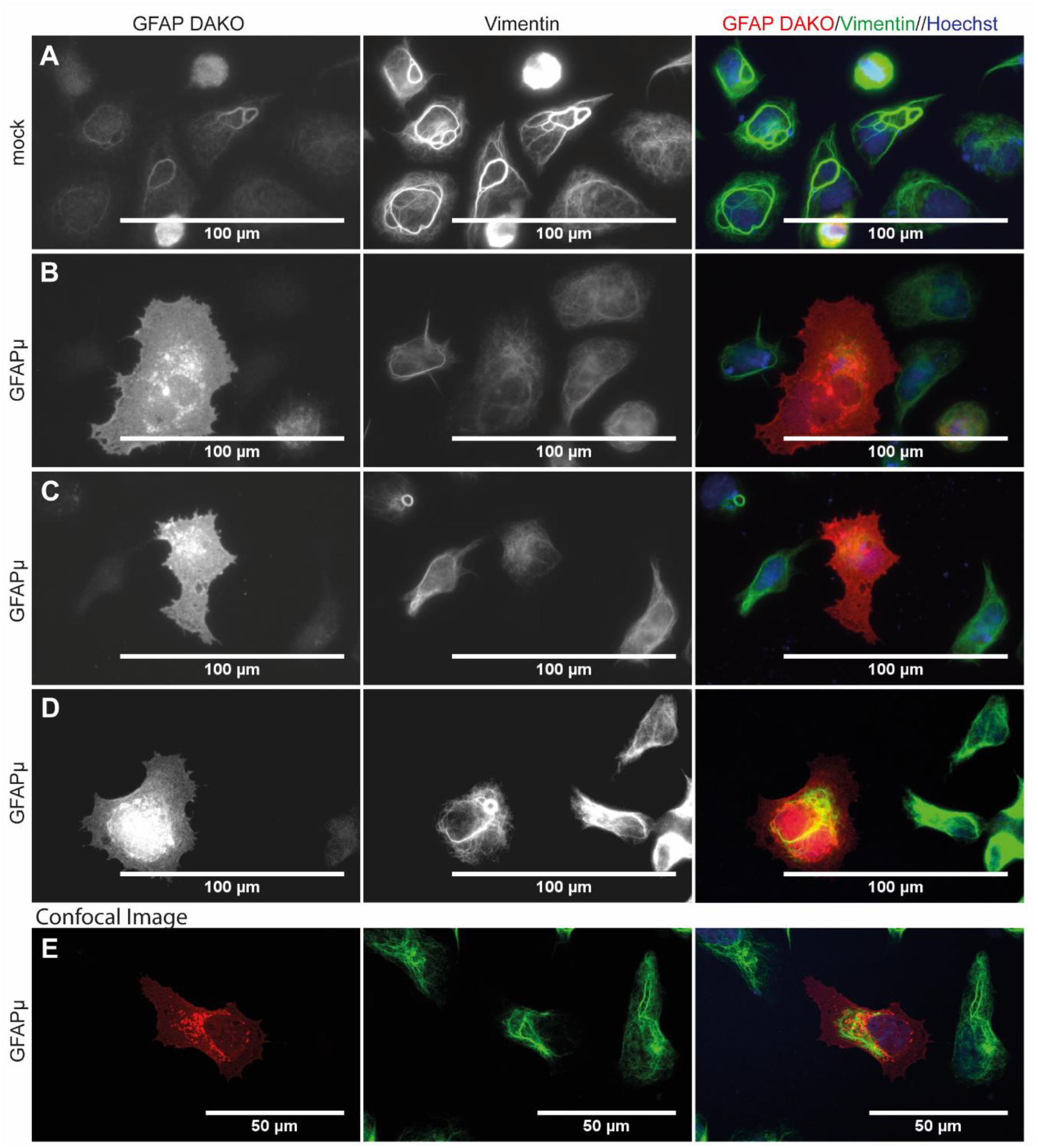
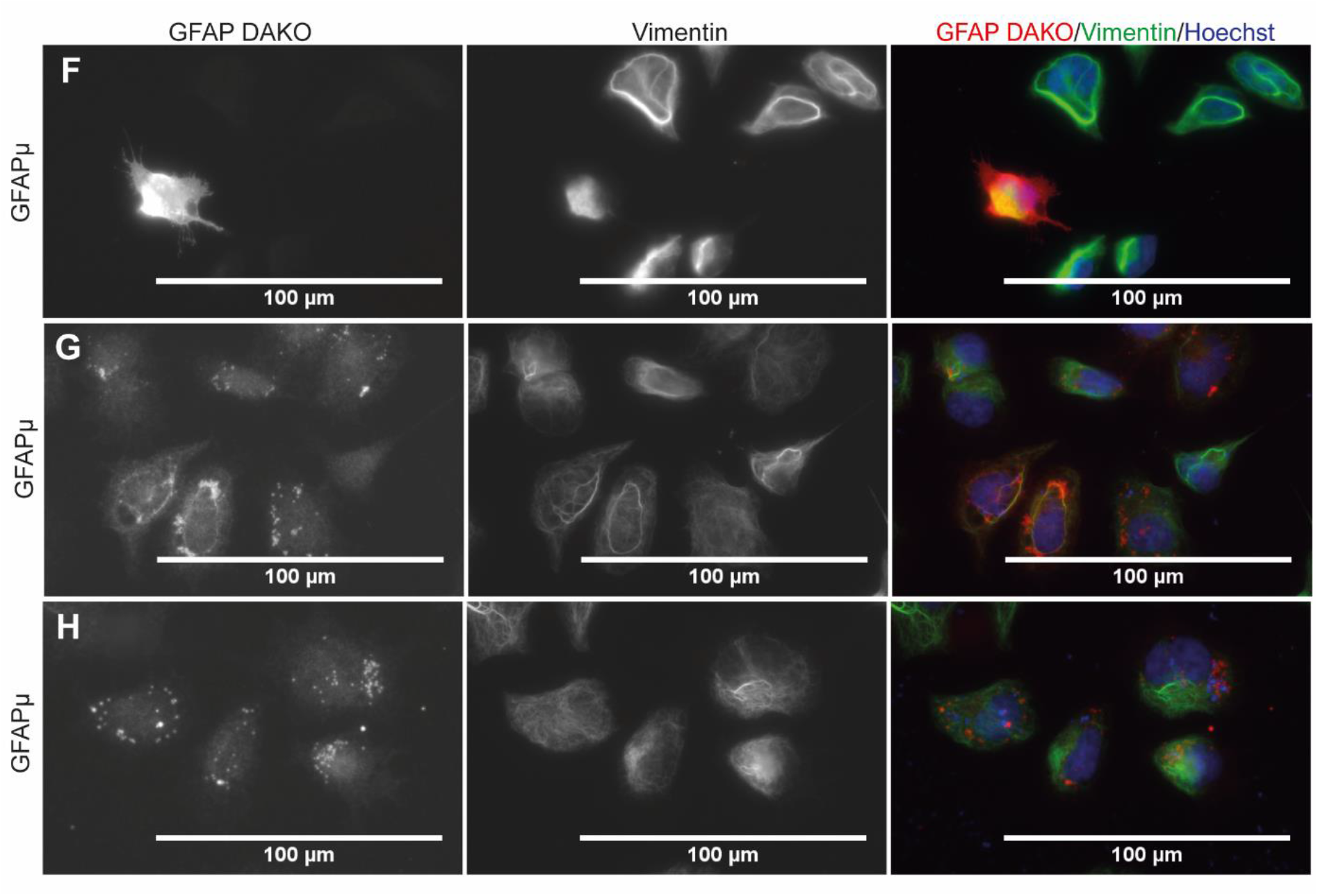
GFAPμ staining patterns in SW13.Vim+ cells. Immunostaining of mock transfected (A) and GFAPμ transfected SW13.Vim+ (B-I) cells for GFAP (Dako) and vimentin. **A** Mock transfected cells. Vimentin staining showed bright and thick filament bundles as well as thinner filamentous, less bright, structures. Some cross-reactivity of the GFAK Dako antibody is observed. **B, C** Cells with diffuse GFAP staining and small (B) or big peri-nuclear (C) aggregates and localization of vimentin thin filamentous structures to the nucleus. **D** Cells with diffuse GFAP immunostaining including large aggregation near the cell’s nucleus. In these cells, vimentin localized to the cell’s nucleus in thick and bright filament bundles. **E** Confocal image of a cell with GFAP positive aggregates that localize to the vimentin intact network. **F** A cell with a large GFAP and vimentin positive peri-nuclear aggregate. **G, H** Cells in which small GFAP aggregates are present that seemed to align alongside bright vimentin filamentous structures.

### GFAPμ expression disrupt the endogenous GFAP network in glioma cells

To determine effect of GFAPμ expression on the GFAP-containing IF network of glioma, we expressed GFAPμ in the U251-MG glioma cells (Figure 5). These cells express endogenous GFAP, vimentin, nestin, and synemin (Moeton et al., 2014). Small and bright GFAP positive aggregates were observed upon GFAPμ expression (Figure 5B, C). Similar to the observations in SW13.Vim+ cells (Figure 4), the aggregates did not contain vimentin indicating that GFAPμ does not affect vimentin filaments.

**Figure 5.**
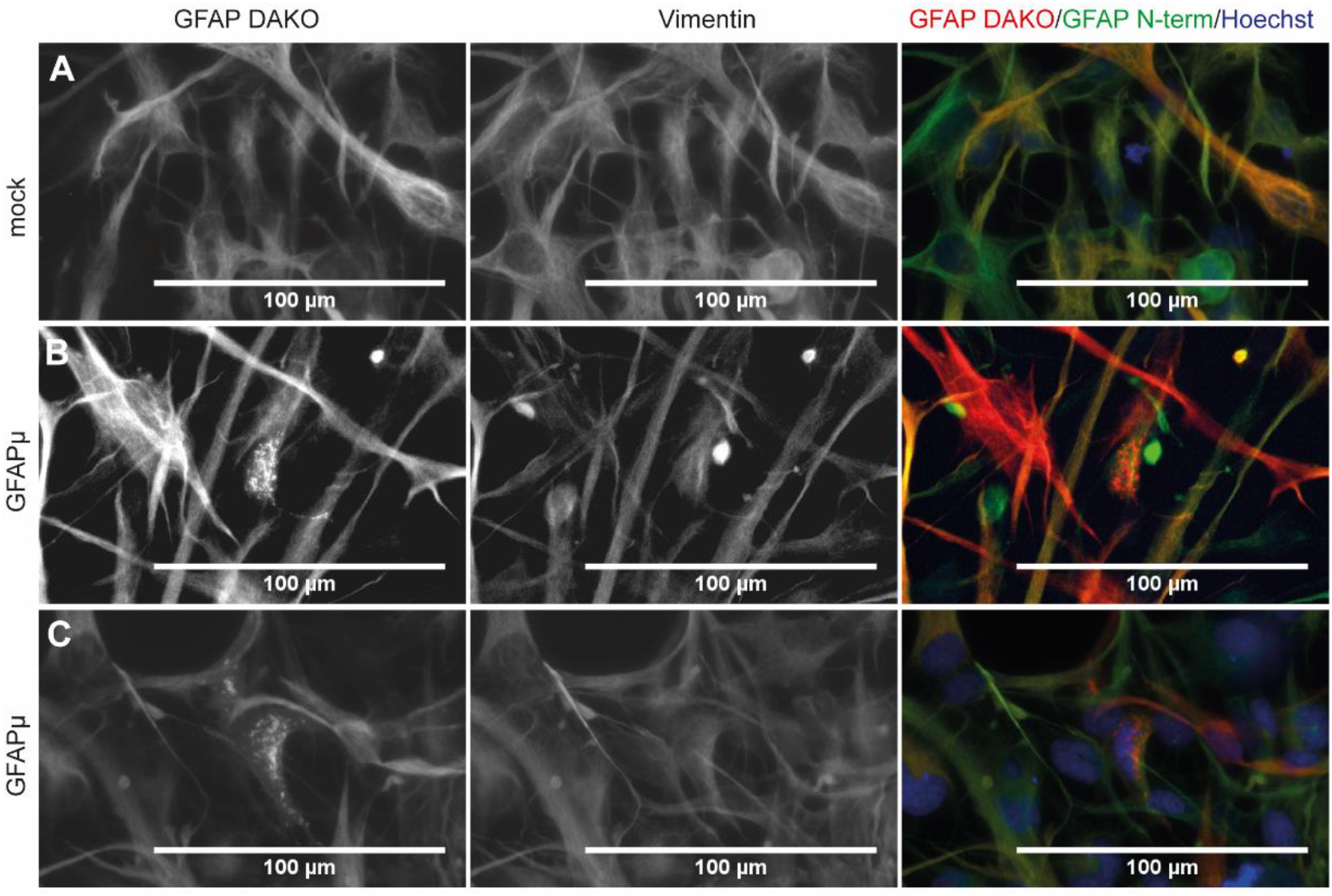
GFAPμ staining patterns in U251-MG cells. Immunostainings of mock (A) and GFAPμ (B, C) transfected U251-MG cells. U251-MG cells were stained for GFAP Dako (left), vimentin (middle), and Hoechst. **A** Immunostaining of U251-MG mock transfected cells showed a characteristic GFAP and vimentin intermediate filament network. **B, C** GFAPμ transfected cells contained GFAP positive small and bright aggregates that are negative for vimentin.

The GFAP antibody used here recognizes all GFAP isoforms, including GFAPμ (Figure 5). Therefore, GFAPμ protein was biotinylated to asses the effect of GFAPμ on the endogenous GFAP network of glioma cells without the availability of a GFAPμ-specific antibody. We generated a construct that encodes GFAPμ protein with an N-terminal 12 amino acid long tag (bio-GFAPμ) that is recognized and biotinylated by the E. coli biotin ligase BirA. Streptavidin labelling of cells co-transfected with BirA and bio-GFAPμ specifically visualized bio-GFAPμ protein. Co-localization analysis of streptavidin and GFAP immunostaining indicated where and how GFAPμ protein is expressed in the cell.

Bio-GFAPμ aggregated in all bio-GFAPμ expressing U251-MG cells (Figure 6). Intense streptavidin labelling of aggregates in bio-GFAPμ and GFP co-transfected cells (Figure 6A) and GFAP positive aggregates in bio-GFAPμ transfected cells (Figure 6C-F) showed that the GFAP positive aggregates contain GFAPμ protein. Confocal imaging showed the presence of GFAP positive and streptavidin negative aggregates as well, indicating that GFAPμ disrupts the endogenous GFAP network (Figure 6E, F, indicated with arrows). To exclude an effect of N-terminal biotin on the assembly of GFAP IFs, bio-GFAPα and BirA were co-expressed in SW13.Vim-cells. Bright streptavidin fluorescence of GFAP filaments similar to filaments observed upon the expression of unlabeled GFAPα were observed (Supplementary Material 4). We therefore excluded the possibility of a sole effect of biotin to the disruption of the endogenous GFAP network and attribute our observations to the expression of GFAPμ.

**Figure 6.**
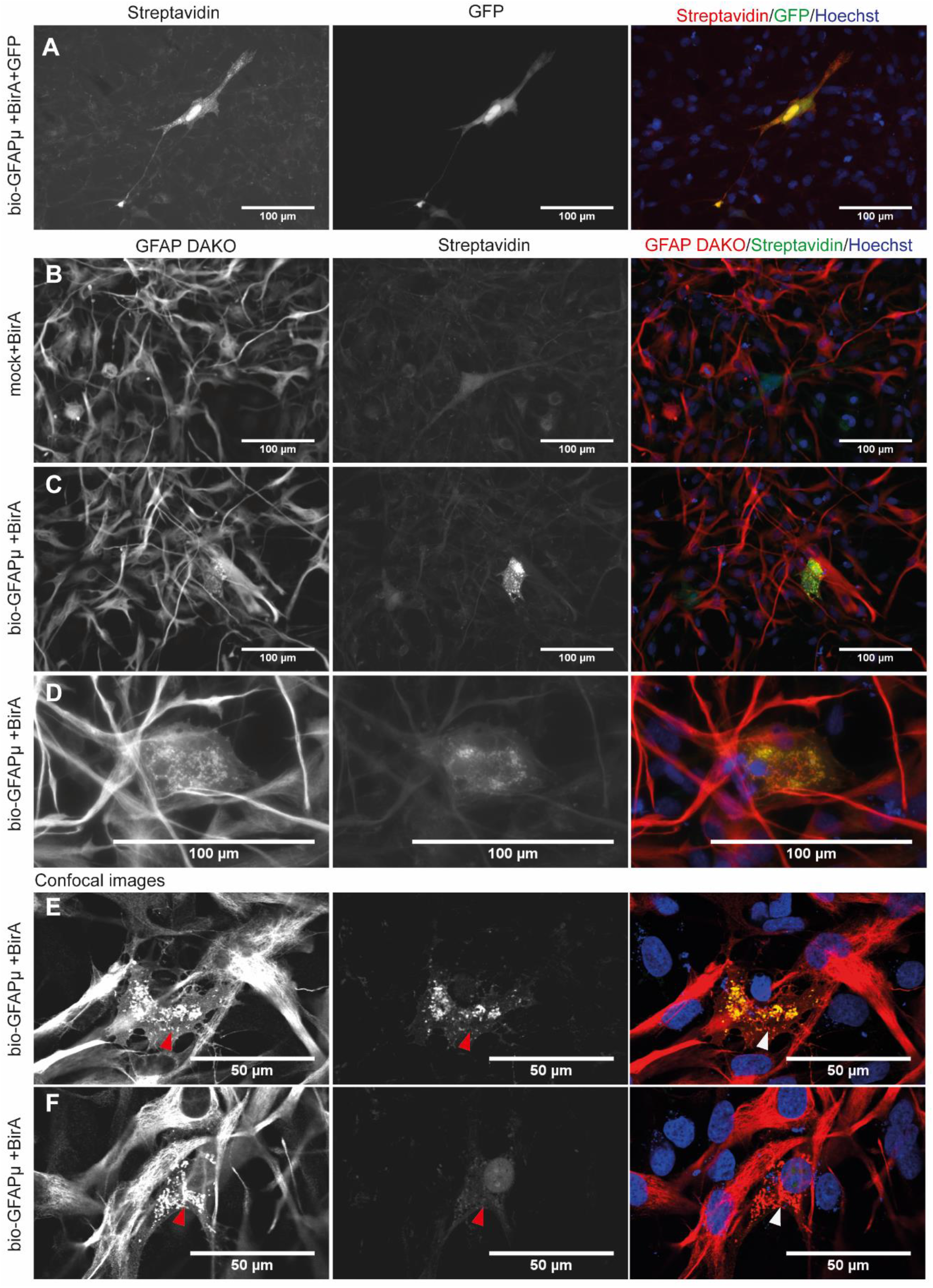
Visualization of bio-GFAPμ in the GFAPμ-induced disrupted U251-MG IF network. Immunostaining of U251-MG cells for GFAP. Biotinylated proteins were labelled with streptavidin. The nucleus was counterstained using Hoechst. **A** Cells transfected with bio-GFAPμ, BirA and GFP (transfection control) and labelled for streptavidin. Streptavidin positive GFAPμ characteristic aggregates are found in the GFP positive cells. **B** Control cells in which empty pcDNA3.1 was co-transfected with BirA to determine background biotin and biotinylation. **C, D** GFAP immunostaining and streptavidin labelling of cells co-transfected with bio-GFAPμ and BirA show co-localization of GFAP and streptavidin in bright streptavidin positive cells. **E, F** Confocal images of cells co-transfected with bio-GFAPμ and BirA that show GFAP positive and streptavidin negative (arrows) aggregates.

## Discussion

The process of alternative splicing is essential for the diversification of cellular phenotype and function. Differences in RNA isoform expression between tissue types and alterations in various types of cancers have been identified by RNA sequencing (Elkon et al., 2013; Bentley, 2014; Herzel et al., 2017). In glioma, alterations in the expression of the type III IF GFAP isoforms GFAPα and GFAPδ have functional implications for the malignant behavior of tumor cells and discrimination between GFAP isoforms could benefit glioma diagnostics (Moeton et al., 2014, 2016; Stassen et al., 2017; van Bodegraven et al., 2019a; b). In this study, the analysis of RNA sequencing data of glioma patient material has revealed the expression of another and new GFAP isoform in glioma, GFAPμ. The GFAPμ transcript was detected in different glioma subtypes by RNA sequencing and was significantly decreased in grade IV glioma. The expression of GFAPμ mRNA, a product of GFAP exon 2 skipping which results in a PTC in exon 3, was confirmed by qPCR analysis in healthy brain tissue, glioma cell lines, and primary glioma cells. We here provide evidence for a new GFAP alternative splicing event observed in glioma and in the healthy brain that results in the expression of GFAPμ.

The detection of a GFAPμ specific peptide in a human proteomics study (Kim et al., 2014) and a ^~^21 kDa GFAP protein in the intermediate filament fraction of proteins isolated from the human spinal cord in this study, provide evidence for translation of the transcript and endogenous expression of GFAPμ protein. Despite these observations, it is important to note that the GFAPμ transcript might be targeted by the process of nonsense-mediated RNA decay (Hug et al., 2016). As GFAPμ contains a PTC in close proximity to an exon-exon junction (51 bp for GFAPμ), it might be recognized as an error during the translation process and being targeted for degradation, as was predicted for this transcript in the Ensembl database (GFAP-202, www.ensembl.org). As most PTC-bearing mRNAs are downregulated by the enhancement of splicing of the canonical transcript (Gudikote et al., 2005), and GFAPμ was detected at ^~^5000 times lower levels compared to GFAPα in RNA sequencing data of TCGA, this possibility should not be neglected. However, as evidence supports endogenous expression of GFAPμ protein, it is more likely that an alternative polyadenylation signal within the 3’UTR of GFAPμ exon 3 is recognized resulting in removal of the exon 3 to exon 4 junction from the transcript and the stable translation of GFAPμ.

We further showed that inducing the expression of the GFAPμ CDS generated a ^~^21 kDa protein. This short GFAPμ protein behaved as expected from its sequence and had a low capacity to self-assemble and formed aggregates in the absence of other GFAP proteins. We observed some short and thick filament-like structures in the absence of other cytoplasmic IFs that could be evidence of filament precursors called squiggles (Chou et al., 2007). These structures form at the cell periphery and when they fail to integrate into a network they aggregate. In glioma cells, the induced expression of GFAPμ disrupted the existing GFAP IF network. The presence of GFAP positive and biotin-negative small and bright aggregates, showed that GFAPμ can induce aggregation of the endogenous GFAP network. Induced overexpression of the self-assembly incompetent GFAPδ isoform in glioma cells disrupts the IF network of glioma cells as well (Moeton et al., 2016). However, increased endogenous GFAPδ protein leaves the IF network intact and impacts glioma cell malignant behavior (van Bodegraven et al., 2019b). In these cells, higher levels of GFAPδ are observed within the IF network. As GFAPμ is expressed 5,000 times lower compared to GFAPα in glioma, the induced expression levels of GFAPμ in our experiments far exceed the endogenous level. We therefore hypothesize that, like GFAPδ, GFAPμ might integrate into the IF network when expressed at endogenous levels and thereby contribute to the function of glioma cells.

Glioma consist of a heterogeneous population of GFAP positive cells (van Bodegraven et al., 2019a). We previously described that the increased relative expression of GFAPδ to GFAPα might distinguish a GFAP positive subpopulation of high malignant glioma cells with invasive characteristics (Stassen et al., 2017; van Bodegraven et al., 2019b; a). As GFAPμ expression was decreased in high malignant glioma and absent in GFAP expressing ihNSCs and adult neural stem cells, we hypothesize that GFAPμ is expressed in lower malignant glioma and more differentiated cell types. Future studies that analyze GFAPμ expression in different types of tissue and cells and determine the functional consequences of GFAPμ expression and integration into the IF network, are needed to confirm this hypothesis.

The here reported differential expression of GFAPμ in glioma subtypes in addition to GFAPα and GFAPδ further emphasizes the importance of GFAP isoform-specific analysis in glioma diagnostics and research. GFAP alternative splicing could form an interesting therapeutic target for glioma treatment. Mechanistic knowledge on GFAP RNA processing and functional consequences of GFAP isoform-specific IF networks for glioma malignancy are needed to further evaluate this possibility.

## Acknowledgements

We would like to thank Vanessa Marques Donega and Marjolein Sneeboer for providing us with RNA samples of primary adult neural stem cells and human brain tissue. The results shown here are in part based on data generated by the TCGA Research Network: http://cancergenome.nih.gov/. This work was supported by the Netherlands Organization for Scientific Research [NWO; VICI grant 865.09.003], the T&P Bohnenn fund, the Dutch Cancer Society [KWF 10123] and the Netherlands Brain Bank (NBB), which is supported by the Netherlands Organization for Scientific Research (NWO).

## Data availability

The results in this study are in part based on data generated by the TCGA Research Network: http://cancergenome.nih.gov/ and the Ensembl database: www.ensembl.org. All other data is available on request from the authors.

---

**Supplementary Material 1.**
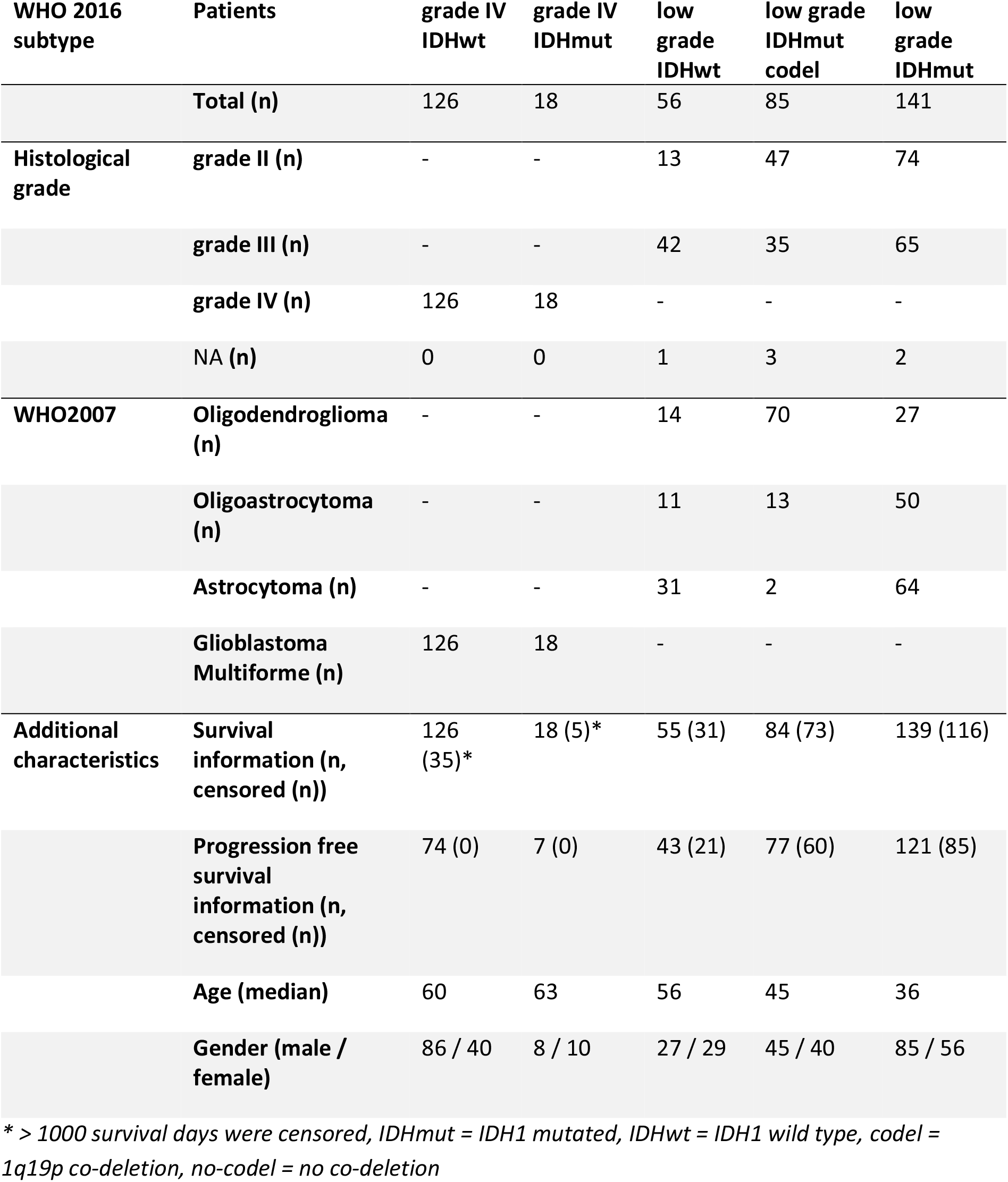
Clinicopathological characteristics of patients of The Cancer Genome Atlas included in the RNA sequencing data analysis

**Supplementary Material 2.**
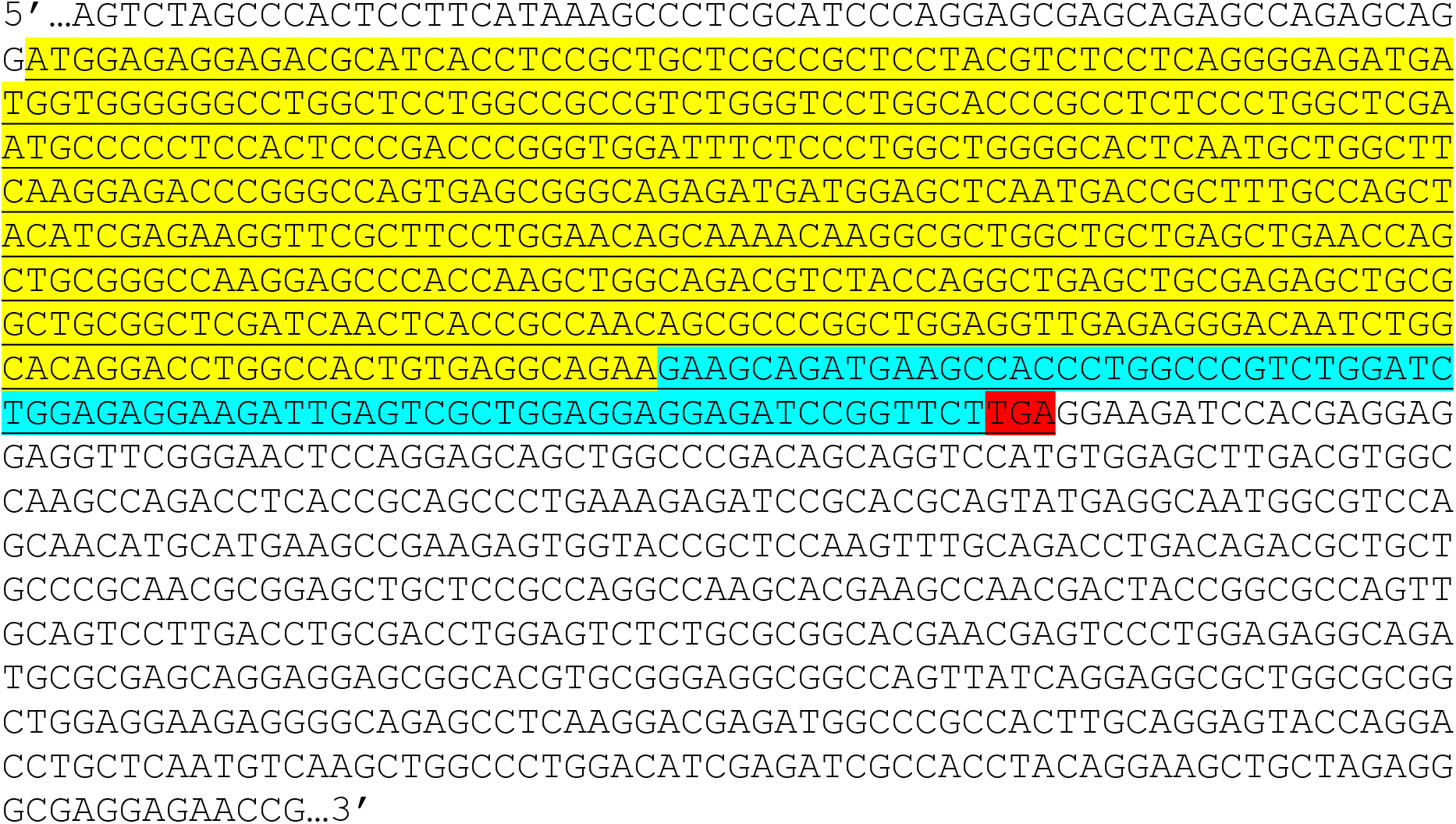
Transcript sequence of GFAPμ. The sequence was used for alignment of RNA sequencing data from astrocytoma grade II, III and IV patients (uc010wjg.1, The Cancer Genome Atlas) (underlined = coding sequence (CDS), yellow = CDS exon 1, blue = CDS exon 3, red = premature termination codon (PTC)).

**Supplementary Material 3.**
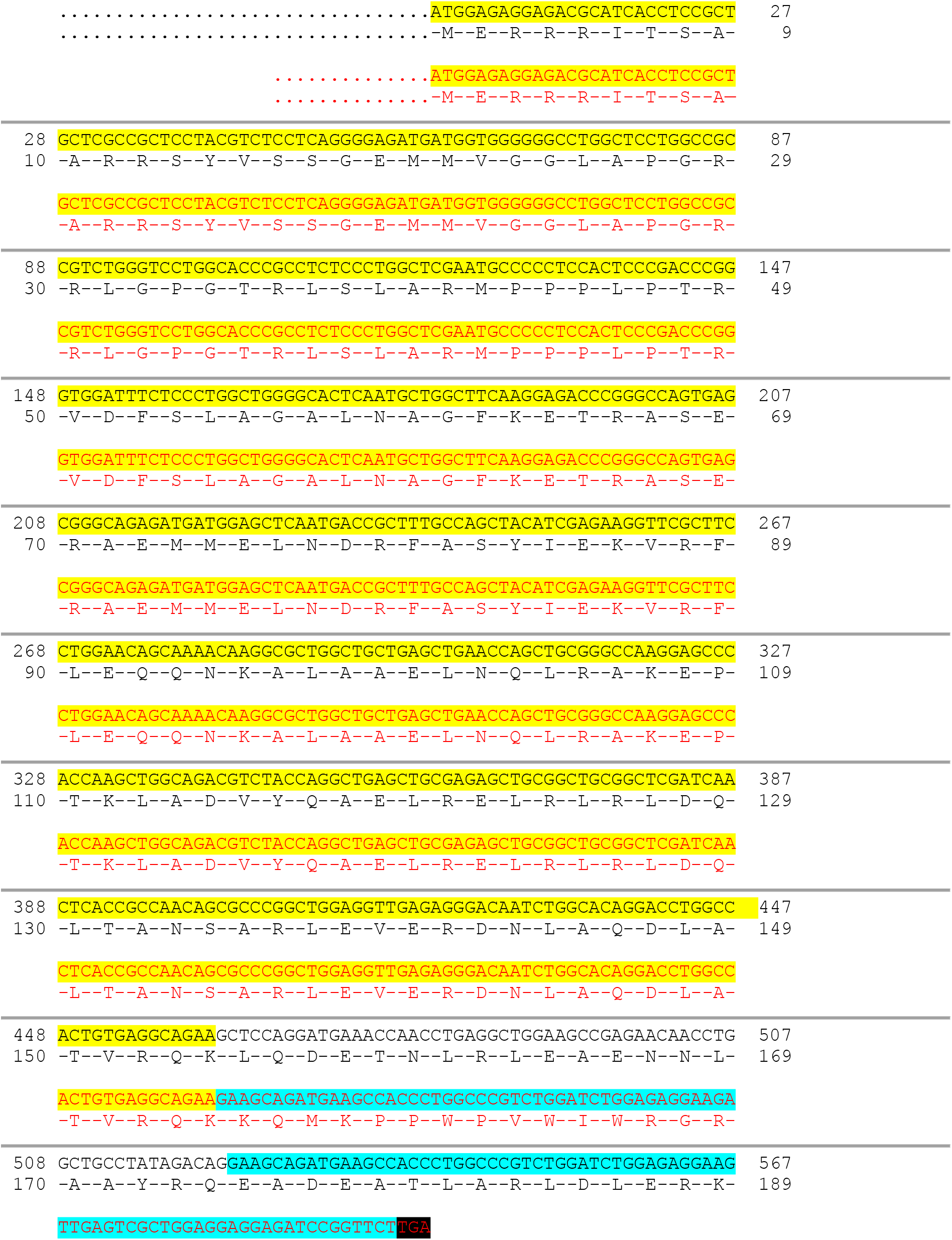
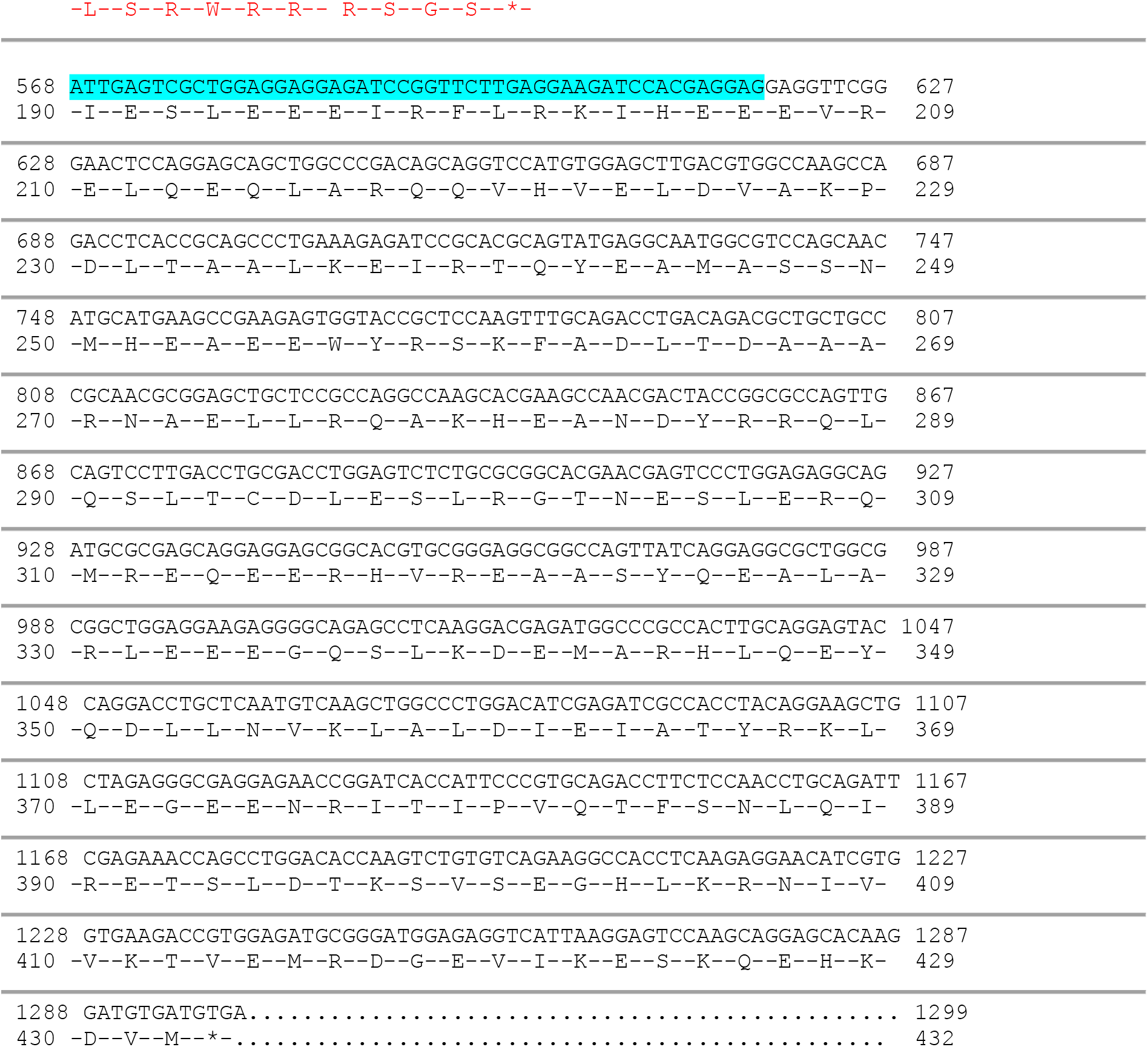
GFAPα and GFAPμ coding and protein sequences. Coding sequence and protein sequence of the canonical GFAP isoform (GFAPα, black) compared to GFAPμ (red) (exon 1 = yellow, exon 3 = blue, PTC = black).

**Supplementary Material 4.**
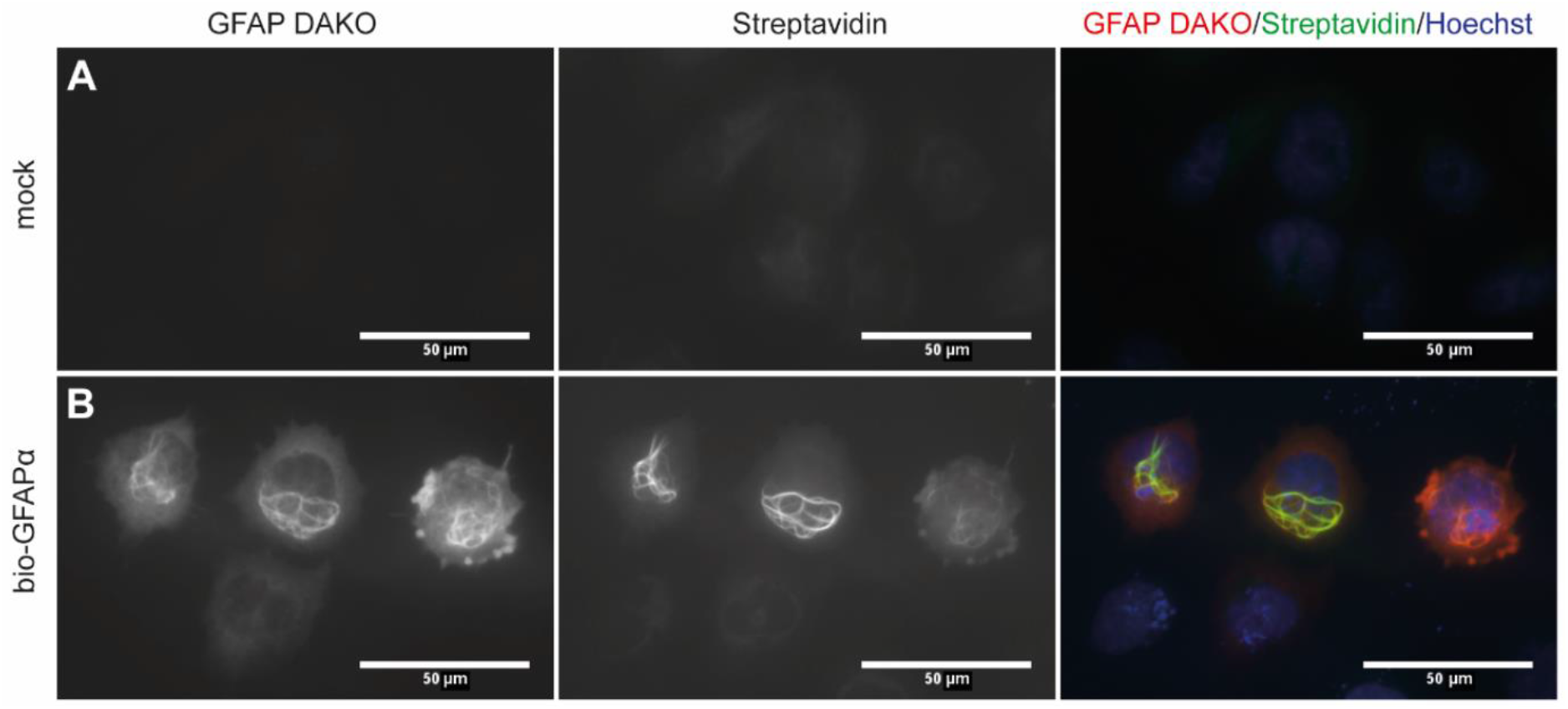
Bio-GFAPα network formation in SW13.Vim-cells. Immunostaining of SW13.Vim-cells for GFAP. Biotinylated proteins were labelled with streptavidin. The nucleus was counterstained using Hoechst. **A** Immunostaining of mock (BirA only) transfected cells. Some background streptavidin immunostaining of endogenous biotin, or aspecific biotinylation of endogenous proteins by BirA is observed. **B** Immunostaining of cells transfected with bio-GFAPα and BirA with bright streptavidin positive filament bundles that co-localized with GFAP immunostaining.

